# TECPR1 provides E3-ligase like activity to the ATG5-ATG12 complex to conjugate LC3/ATG8 to damaged lysosomes

**DOI:** 10.1101/2023.06.24.546289

**Authors:** Yingxue Wang, Matthew Jefferson, Matthew Whelband, Kristine Kreuzer, James Mccoll, Paul Verkade, Ulrike Mayer, Thomas Wileman

## Abstract

Autophagosomes deliver cytosolic material to lysosomes to provide amino acids during starvation and to degrade damaged proteins and organelles to maintain tissue homeostasis. Delivery to lysosomes requires LC3/ATG8 (LC3), the major membrane protein of the autophagosome, that uses adaptor proteins to capture cargo and recruits tethering and SNARE proteins to promote fusion with lysosomes. LC3 is also recruited to endo-lysosome compartments in response to increases in vacuolar pH to facilitate degradation of material entering cells by endocytosis. A series of ubiquitin-like reactions conjugate LC3 to amino groups exposed by phosphatidylethanolamine (PE) or phosphatidylserine (PS) in target membranes. The E1 and E2-ubiquitin-like activities of ATG7 and ATG3 use thioester bonds to transfer LC3 to the ATG5-ATG12 conjugate. At the same time binding of ATG16L1 to ATG5-ATG12 provides the E3 ubiquitin-ligase like activity necessary to conjugate LC3 to PE or PS. The ATG5-ATG12 conjugate can also bind TECPR1 (tectonin beta propeller repeat-containing protein) which shares an ATG5 interaction (AIR) region with ATG16L1 and can binds LC3 directly through a LC3 interaction region (LIR). In this study we have used cells lacking ATG16L1 to determine if TECPR1 can substitute for ATG16L1 during LC3 conjugation. The results show that ATG16L1-/-MEFS can conjugate LC3 to lysosomes damaged by chloroquine or LLOMes and conjugation is dependent on the ubiquitin-like enzymes ATG3, ATG5 and ATG7 upstream of ATG16L1. TECPR1, ATG5 and galectin 3 are recruited to LAMP positive damaged lysosomes in the absence of ATG16L1 suggesting that TECPR-1 recruits ATG5-ATG12 to conjugate LC3 to damaged lysosomes. This was confirmed when truncation of TECPR1 at the central PH domain required for lysosome binding prevented LC3 conjugation, and LC3 conjugation could be restored by full length TECPR1. Recruitment of TECPR1 to damaged lysosomes required the N-terminal LIR motif and was partially dependent on the central PH domain that binds PI4P exposed during lysosome repair. TECPR1 can therefore conjugate LC3 to damaged lysosomes independently of ATG16L1 by providing E3 ligase-like activity to ATG5-ATG12. Direct conjugation of LC3 by TECPR1 may contribute to the autophagosome tethering functions reported for TECPR1 by increasing recruitment of cargo receptors, tethering proteins and SNARE proteins required for fusion with lysosomes and protein degradation. TECPR1-dependent conjugation of LC3 may also facilitate lysosome repair pathways involving autophagic lysosomal reformation and transcriptional activation of autophagy by TFEB.

## Introduction

Autophagy is a highly conserved membrane trafficking pathway that delivers cytosolic material to lysosomes for degradation [1-3]. Autophagy can provide a short-term supply of amino acids during starvation and can degrade damaged proteins and organelles to maintain tissue homeostasis [1, 2, 4-6]. Delivery to lysosomes is mediated by several conserved autophagy (ATG) proteins which generate autophagosomes to engulf material in the cytosol and then fuse with lysosomes [7-9]. LC3/ATG8 (LC3) is the major membrane protein of the autophagosome [10]. LC3 on the inner surface of the expanding autophagosome facilitates uptake of cargos by binding adaptor proteins such as p62 [11, 12]. Once autophagosomes have sealed, LC3 exposed to the cytosol binds microtubule motor proteins to transport autophagosomes towards lysosomes where LC3 provides a platform for interactions with tethering proteins (HOPS, PLEKHM1) and SNARE proteins (STX17, SNAP29, VAMP8) that promote lysosome fusion and degradation of autophagy cargos by lysosomal proteases [13, 14]. LC3 is also recruited to the cytosolic face of endo-lysosome compartments in response to changes in vacuolar pH induced during the uptake of pathogens, apoptotic cells or transfection agents, or following incubation with lysosomotropic drugs, inhibitors of the vacuolar ATPase (vATPase) or stimulation of lysosome Ca^2+^ channel TRPML1 [15, 16]. These alternative pathways have been called LAP (LC3 associated phagocytosis) or CASM (<u>c</u>onjugation of <u>A</u>TG8 to <u>s</u>ingle <u>m</u>embranes) and are thought to facilitate degradation of endocytic cargoes in lysosomes [16-19].

Early studies showed that a subset of core autophagy proteins provide ubiquitin-like enzyme reactions to conjugate LC3 to amino groups exposed by phosphatidylethanolamine (PE) in the autophagosome membrane [20-23]. This ubiquitin-like pathway starts with processing of LC3 by the ATG4 cysteine protease to expose a C-terminal glycine residue [24]. The E1-ubiquitin-like activity of ATG7 then generates a thioester intermediate between an active site cysteine and the C-terminus of LC3 [21]. Subsequent hydrolysis of ATP allows an E2-like reaction to transfer LC3 to the active cysteine of ATG3 generating an ATG3-LC3 ‘E2-carrier’ complex [22]. The last stage in transfer of LC3 to PE requires the E3 ubiquitin-ligase like activity of the ATG5-ATG12:ATG16L1 complex where ATG5 induces a conformational change in the catalytic site of ATG3 leading to transfer of the C-terminal carboxyl group of LC3 from a thioester bond to amino groups exposed by PE [25]. Correct delivery of LC3 to the autophagosome membrane is mediated by ATG16L1 within a large ∼800kDa complex containing hexamers of ATG5-ATG12:ATG16L1 [26, 27]. In common with ubiquitin E3 ligases which direct ubiquitination to specific substrates, ATG16L1 determines the site of conjugation of LC3 to membranes [26]. The coiled-coil domain of ATG16L1 binds WIPI2, which is recruited to PIP3 (phophatidylinositol-3 phosphate) generated early during autophagy by VPS34 within the PI3KC3 (class III phosphatidylinositol 3-kinase) complex [28]. This leads to recruitment of multiple copies of the ATG16L1:ATG5-ATG12 complex to sites of autophagosome expansion, where activation of the ATG3-LC3 intermediate by ATG5-ATG12 conjugates LC3 to PE. The ubiquitin-like conjugation reactions of ATG7, ATG3 and the ATG5-ATG12:ATG16L1 complex are also required for conjugation of LC3 to endolysosome compartments [29]. In this case the WD domain of ATG16L1 binds to the vATPase in the vacuolar membrane leading to conjugation of LC3 to PE or phosphatidylserine (PS) [17, 30].

Proteomic network analysis [31] yeast two hybrid screens and pull downs [32, 33] have shown that the ATG5-ATG12 conjugate can also bind TECPR1 (tectonin beta propeller repeat-containing protein). A comparison of the crystal structures of ATG16L1:ATG5 and TECPR1:ATG5 complexes [34] shows that a-helices in ATG16L1 and TECPR1 share a common ATG5 interaction (AIR) region that binds ATG5. TECPR1 also binds LC3 directly through a LC3 interaction region (LIR) motif and a central PH domain allows TECPR1 to bind phospholipids [33, 35]. Recruitment of the TECPR1:ATG5-ATG12 complex is thought to tether autophagosomes to lysosomes to facilitate autophagosome-lysosome fusion [34], mitophagy and selective autophagy of bacterial pathogens [32]. Crystal structures of the ATG5 binding domains of TECPR1 and ATG16L1 show that the a-helical <u>A</u>TG<u>5 i</u>nteraction <u>m</u>otif (AFIM) **W-**x-x-x-**I**-x-x-**L**-x-x-**R**-x-x-x-**Q/E** of TECPR1 can be superimposed on the AFIM of ATG16L1 where both align along the b-strands the ubiquitin-fold domains of ATG5 [34]. The observations that the TECPR1 binds LC3 and that the TECPR1 AFIM occupies the same space within the ubiquitin fold domain of ATG5 as ATG16L1 raised the possibility that TECPR1 could substitute for ATG16L1 and form TECPR1:ATG5-ATG12 complex able to conjugate LC3 to PE or PS independently of ATG16L1. This study has tested this possibility by incubating cells lacking ATG16L1 under conditions known to activate autophagy. The results show that LC3 is conjugated to damaged lysosomes independently of ATG16L1 by a pathway that is dependent on TECPR1 which provide E3 ligase-like activity to ATG5-ATG12. This may contribute to the tethering functions of TECPR1 reported promote autophagosome lysosome fusion and/or facilitate recruitment of proteins involved in lysosome repair.

## Results

### Lipidation of ATG8/LC3 can occur independently of ATG16L1

MEFS lacking expression of ATG16L1 (ATG16L1-/-) were incubated under conditions known to stimulate autophagy-related pathways that lead to translocation of LC3 from the cytoplasm to membranes. Figure 1A shows that WT MEFS generate LC3 puncta after induction of autophagy by starvation in HBSS (ii) and generate large LC3 positive vacuoles following induction of non-canonical autophagy/LAP/CASM by chloroquine (iii). Cells lacking ATG16L1 (Figure 1A iv-vi) were unable to generate LC3 puncta in response to HBSS (v) but interestingly the cells generated bright LC3 puncta after incubation with chloroquine (vi). Unlike WT cells, LC3 was not recruited to the ring-like perimeter of swollen vacuoles in the absence of ATG16L1 but was located to bright puncta close to vacuoles swollen by chloroquine. Counter staining showed that these smaller puncta contained lysosome membrane protein LAMP1 (see ROI) and were present in both cell types. Generation of LC3 puncta following incubation with chloroquine was not observed for cells lacking proteins upstream of ATG16L1 in the LC3 conjugation pathway (Figure 1B) such as the ubiquitin-like enzymes ATG3 (i-iii), ATG5 (iv-vi) or ATG7 (vii-ix). The translocation of LC3 to puncta in the absence of ATG16L1 therefore required the E1 like activity of ATG7 and E2-like activity of ATG3 and the ATG5-ATG12 conjugate.

**Figure 1.**
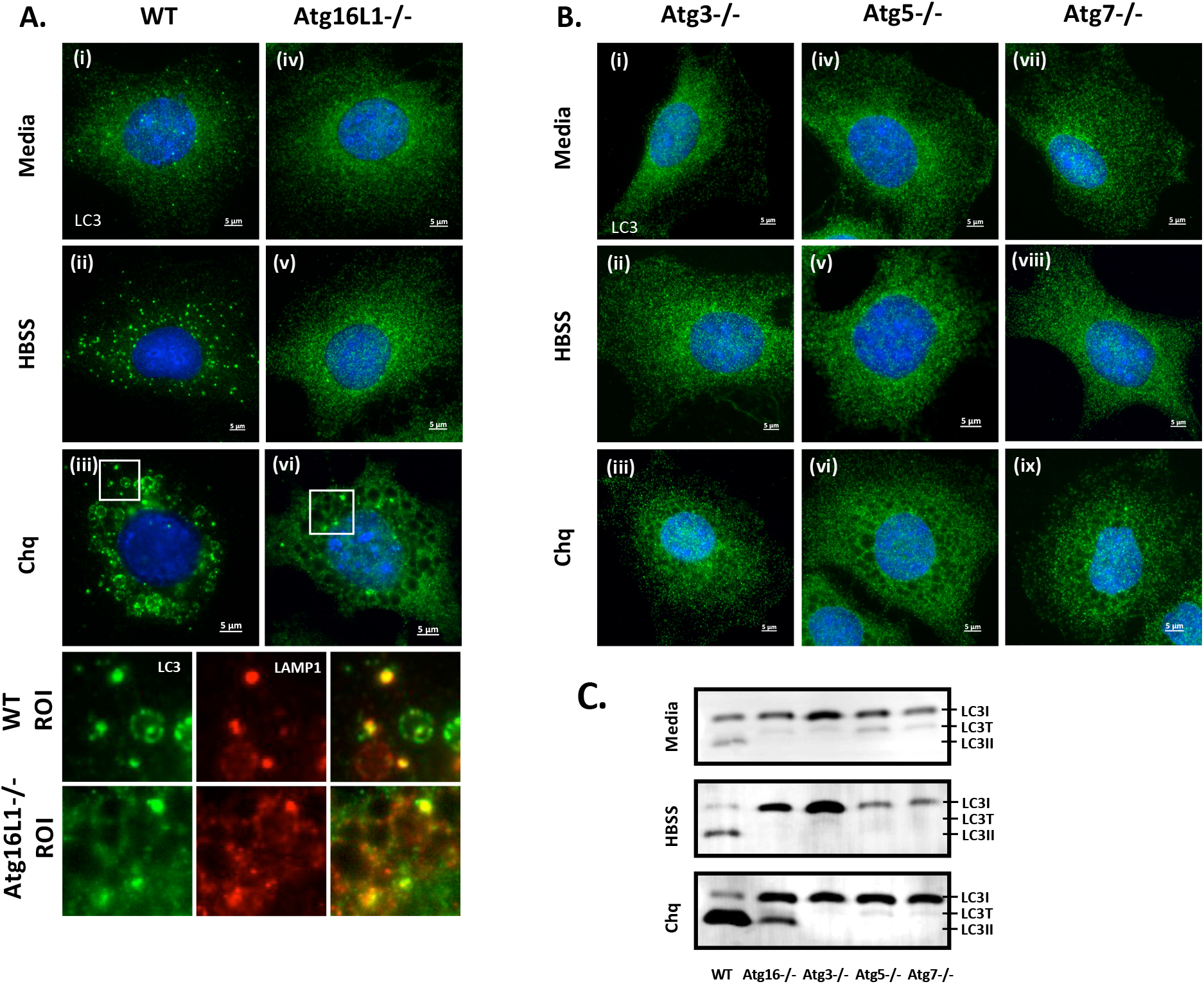
Chloroquine induces formation of LC3 puncta and LC3II independently of ATG16L1. **Panel A**. Atg16L1-/- and control MEFs were incubated for 2 hours in nutrient media (i and iv), HBSS (ii & v) or nutrient media containing chloroquine (100mM iii-vi) as indicated. Cells were fixed and immunostained for LC3. Regions of interest (ROI) from cell incubated with chloroquine are shown at higher magnification with counter staining for LAMP1 (red). **Panel B**. Atg3-/-, Atg5-/- and Atg7 -/- MEFs were incubated for 2 hours in nutrient media as indicated. Cells were fixed and immunostained for LC3. **Panel C**. Cells were incubated as described above and generation of LC3II was assessed by western blot for LC3.

Conjugation of LC3 to membranes involves covalent attachment of LC3 to PE or PS resulting in formation of LC3II which has a characteristic increase in mobility on western blots. Western blots (Figure 1C) show the formation of LC3II in WT cells starved in HBSS or incubated with chloroquine. Close inspection of western blots showed two forms of LC3 migrating faster than LC3I. The slower of these two forms migrating between LC3I and LC3II was likely to be LC3-T where LC3 is partially degraded by the 20S proteasome and cannot be conjugated to PE [36]. Subsequent experiments took care to ensure estimates of LC3II production focused on the faster migrating LC3II and not LC3-T. Figures 1C shows that LC3II was produced by wild type cells incubated in HBSS or chloroquine. LC3II was absent from Atg16L1 -/- cells incubated with HBSS but LC3II was induced by chloroquine. The levels of LC3II were lower than seen in WT cells and this reflected the recruitment of LC3 to small puncta rather than the limiting membrane of large swollen vacuoles. As seen for LC3 puncta, chloroquine was unable to induce LC3II in cells lacking ATG3, ATG5 or ATG7. Taken together the results suggested that chloroquine could activate an ATG16L1-independent pathway downstream of ATG3 and ATG7 that use ATG5-ATG12 to conjugate LC3 to PE or PS.

### ATG16L1 independent lipidation occurs in response to vacuole swelling

Neutralisation of hydrogen ions and consequent osmotic swelling by lysosomotropic drugs such as chloroquine activates CASM/LAP to recruit LC3II to swollen vacuoles. Raised pH can also occur following neuralisation of hydrogen ions by reactive oxygen species (ROS) generated by NADPH oxidase. A panel of inhibitors were used to explore if raised pH or osmotic swelling were responsible for recruitment of LC3 to membranes in Atg16L1-/-cells. The cells were counterstained for LAMP to identify endo-lysosome compartments and images were used to assess co-localisation (Figure 2 A&B) and the corresponding western blots showing formation of LC3II are in figure 2C. Chloroquine generated between 7 and 15 LC3 puncta per cell in Atg16L1^-/-^ MEFS and the majority (90%) co-localised with LAMP (i-iii). The formation of LC3 puncta, and appearance of LC3II on western blot, were not inhibited by wortmannin (iv-v). LC3 recruitment was not therefore dependent of the PI3 kinase activity of vps34 that is upstream of conventional ATG16L1-dependent LC3 conjugation during autophagy. ATG16L1-independent production of LC3 puncta was however reduced by half when the NADPH oxidase was inhibited by DPI (vii-ix) suggesting a link with ROS production which raises vacuole pH. Production of LC3 puncta in cells (Figure 2A x-xii) and detection of LC3II on western blots (Figure 2C) was also inhibited by phloretin which blocks water channels in endosomes and lysosomes and reduces osmotic swelling. The role of osmotic swelling versus raised pH was dissected further using bafilomycin which raises vacuolar pH by inhibiting the v-ATPase but, unlike chloroquine, reduces osmotic swelling by slowing the import of protons into vacuoles. Bafilomycin inhibited both the formation of LC3 puncta (xiii-xv) and appearance of LC3II on western blots, suggesting that osmotic swelling by chloroquine, rather than raised pH was the signal for generation of LC3 puncta in the absence of ATG16L1.

**Figure 2.**
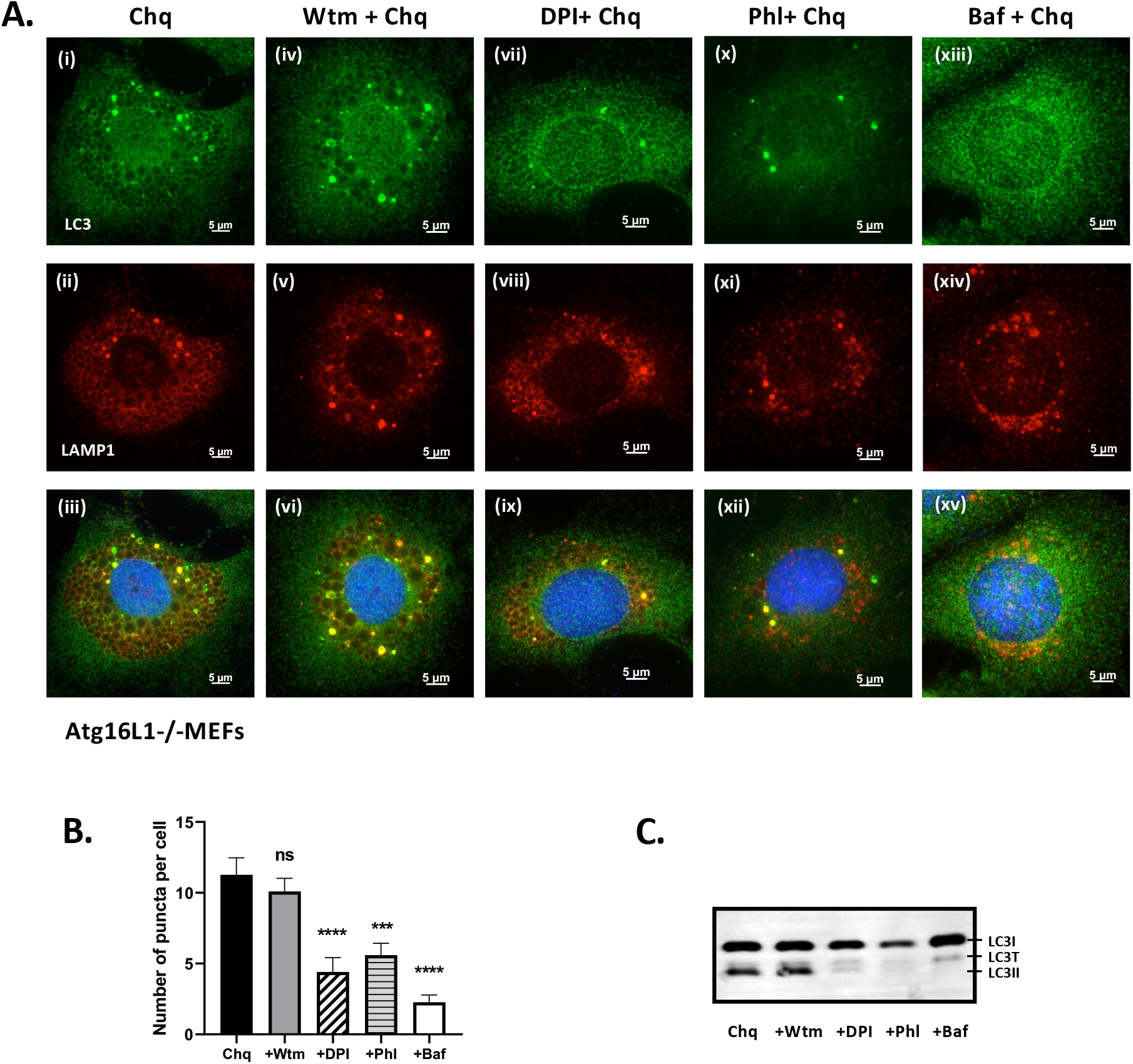
Formation of LC3 puncta and LC3II in Atg16L1-/-cells occurs in response to vacuole swelling. **Panel A**. Atg16L1-/-MEFs were incubated with chloroquine for 2 hours in the presence of wortmannin (100nM iv-vi) DPI (10mM vii-ix), phloretin (180mM x-xii) or bafilomycin (100nm xiii-xv) as indicated. Cells were fixed and immunostained for LC3 (green) and LAMP1 (red). **Panel B**. Numbers of LC3 puncta from more than 10 cells incubated as described in panel A. were quantified by imaris, P-values were calculated using multiple t-test (***P < 0.001). **Panel C**. Cells were incubated as described in panel A and generation of LC3II was assessed by western blot for LC3.

### ATG16L1 independent lipidation facilitates conjugation of LC3 to damaged lysosomes

Osmotic swelling induced by lysosomotropic agents such as chloroquine can lead to damage to vacuoles and exposure of luminal galactoside sugars to the cytosol where they are recognised by galectin-3. The role played by vacuole damage as a signal for conjugation of LC3 to membranes in the absence of ATG16L1 was tested by immunostaining for galectin-3 after incubation of cells with chloroquine (Figure 3A). In control cells (i-iv) bright galectin 3 puncta were seen colocalised with LC3 suggesting accumulation of damaged membranes. These puncta were close to large swollen vacuoles also staining for LC3, but the limiting membrane of the vacuoles was negative for galectin-3 indicating that the swollen vacuoles do not show signs of damage. In ATG16L1-/- cells (Figure 3B i-iv) the galectin 3 again co-localised with bright LC3 puncta, but both galectin-3 and LC3 were absent from the limiting membrane of the swollen vacuoles.

**Figure 3.**
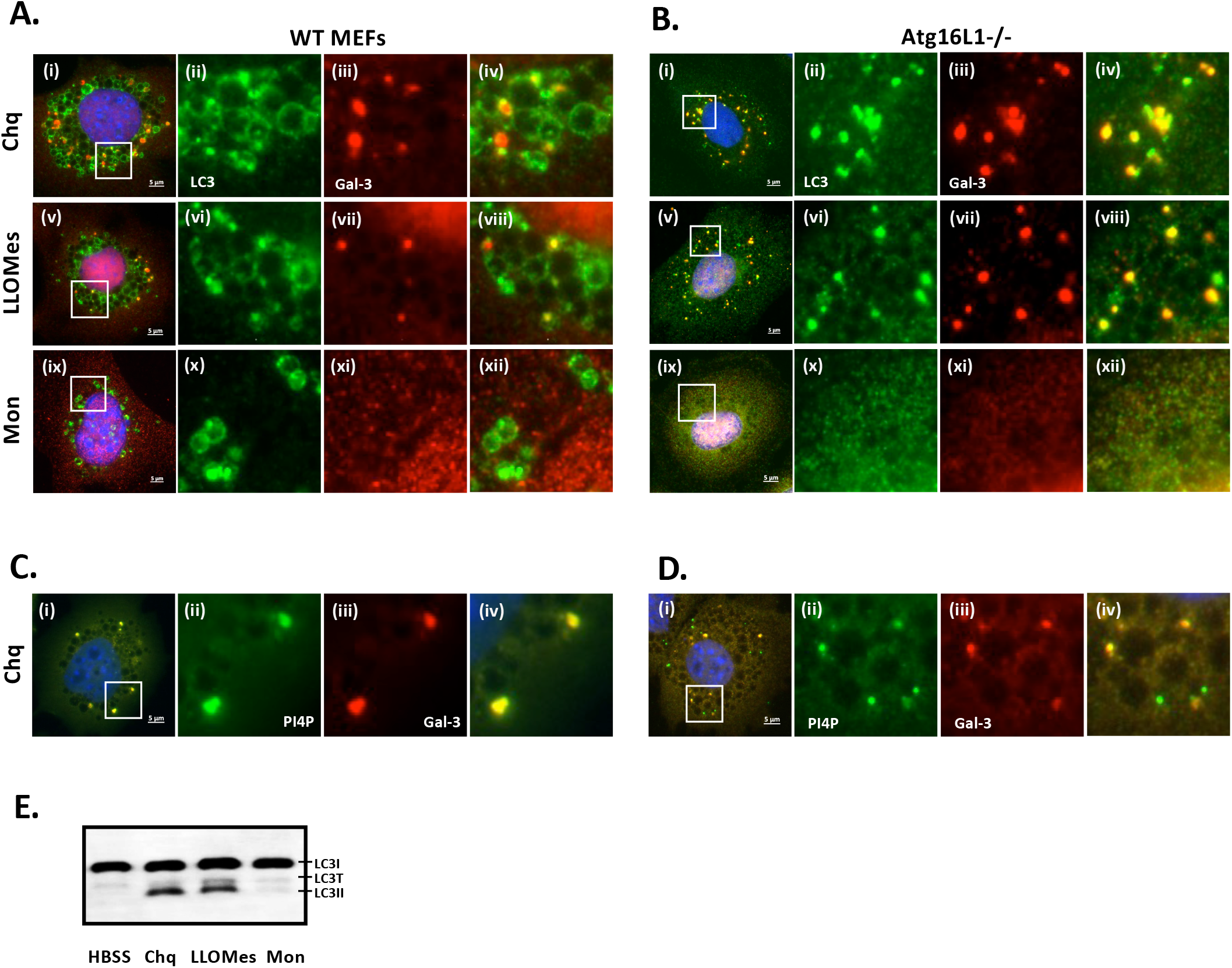
ATG16L1 independent lipidation results in conjugation of LC3 to damaged lysosomes. Control (**Panel A**) and Atg16L1-/-MEFs (**panel B**) were incubated for 2 hours in nutrient media containing chloroquine (100mM i-iv), LLOMes (1mM v-viii) or monensin (100µM ix-xii) as indicated. Cells were fixed and immunostained for LC3 (green) and galectin 3 (red). Control (**panel C**) and Atg16L1-/-MEFs (**panel D**) were incubated for 2 hours in nutrient media containing chloroquine (100mM i-iv) and immunostained for PI4P (green) and galectin-3 (red). **Panel E**. Atg16L1-/-MEFs were incubated as described in panel B and generation of LC3II was assessed by western blot for LC3.

The localisation of lysosome marker LAMP1 in bright LC3 positive puncta generated in response to chloroquine (see figure 1) suggested that the small galectin-3 positive puncta may represent damaged lysosomes. The role played by lysosome damage was tested directly by incubating cells with the lysosome damaging compound Leu-Leu-oMe (LLOMes). In common with chloroquine, LLOMes caused vacuoles to swell. In control cells (Figure 3A v-viii) LLOMes induced recruitment of LC3 to swollen vacuoles (large arrow) and dense puncta (small arrows) and several of the small puncta were positive for galectin 3 (vii). In ATG16L1-/-cells incubated with LLOMes (Figure 3B v-viii) LC3 was absent from swollen vacuoles (v-vi) and was concentrated on small galectin-3 positive puncta (vii-viii). As seen for chloroquine exposure of luminal sugars in response to LLOMes was restricted to small puncta rather than swollen vacuoles. The role played by vacuolar damage was also tested using the ionophore monensin which exchanges protons for Na^+^ and raises vacuolar pH without inducing vacuolar damage. Panels ix-xii of figure 3A and B show that monensin generated swollen vacuoles that recruited LC3 in control cells but not in cells lacking ATG16L1. Monensin failed to generate small puncta positive for galectin-3 or LC3 in either cell type showing that under these conditions monensin does not damage vacuoles. The observation that monensin was unable to generate small LC3 puncta in both cell types coupled with the lack of galectin-3 puncta suggested that vacuolar damage, rather than raised pH is the primary driver for ATG16L1 independent conjugation of LC3 to membranes. This conclusion correlated with western blots (Figure 3E) showing that LC3II was induced in ATG16L1-/-MEFs by chloroquine and LLOMes, but not monensin.

Lysosome damage in response to LLOMes results in recruitment of phosphatidylinositol 4-kinase (PI4K2A) to the lysosome and production of PI4P in the lysosome membrane. The PI4P recruits oxysterol binding protein-related proteins (PRPs) 9-11 to the lysosome allowing interactions with VAPA/B to transport of phospatidylserine (PS) from the ER to the lysosome where it facilitates lysosome repair [37]. Recruitment of PI4P to damaged lysosomes in response to chloroquine was studied by immunostaining for PI4P (Fig 3C and 3D (i-iv)). In agreement with the results above, PI4P was recruited to small puncta staining for galectin-3 in both cell types, but PI4P was not recruited to swollen vacuoles.

### ATG16L1-independent conjugation of LC3 to damaged lysosomes requires TECPR1

Pull down experiments and structural analysis have suggested that the TECPR1 binds ATG5 through an interaction motif shared with ATG16L1 [32-34]. This raised the possibility that a TECPR1:ATG5-ATG12 complex could function in place of ATG16L1 to transfer LC3 from ATG3 to PE/PS in damaged lysosomes. This was tested by using gene editing to inhibit TECPR1 activity in the ATG16L1-/-MEFS (Figure 4A). Guide RNA were introduced into Atg16L1-/-MEFS by lentivirus transduction and cell clones resistant to puromycin were analysed by PCR for disruption of the TECPR1 gene. Sequencing showed that guide RNAs generated a stop codon in exon 12 of the Tecpr-1 gene (supplemental figure 1) disrupting the central PH domain and preventing translation of the C-terminal dysferlin (DysF) and tectonin b-propeller repeat (TECPR). This is labelled TECPR * to indicate truncation rather than complete removal. Figure 4 shows the effect of TECPR1 truncation on LC3 staining of ATG16L1-/-cells. Bright puncta positive for LC3 and galectin-3 were generated in ATG16L1-/-MEFS (Figure 4B i-iii) in response to chloroquine. High magnification (Figure 4B iv) and rendered images (Figure 4B v-vi) showed that puncta were positive for both signals. Galectin-3 positive puncta were also formed in ATG16L1-/-MEFs expressing the truncated TECPR1 (Fig 4B vii-ix) but high magnification (Figure 4B x) and rendered images (xi-xii) showed that these galectin-3 positive puncta were negative for LC3. Quantitation of puncta (Fig 4D) showed an almost complete loss of LC3 puncta following truncation of TECPR1. This was consistent with western blot analysis (Figure 4E) showing that chloroquine was unable to induce LC3II in ATG16L1-/-MEFS following truncation TECPR-1. The experiment was repeated for cells incubated with LLOMes (Figure 4C). Again, galectin-3 puncta were produced in both cell types indicating damaged lysosomes, but damaged lysosomes did not recruit LC3 in cells expressing the truncated TECPR-1. This was confirmed by quantitation of puncta in cells (Figure 4F) and western blot (Figure 4G)

**Figure 4.**
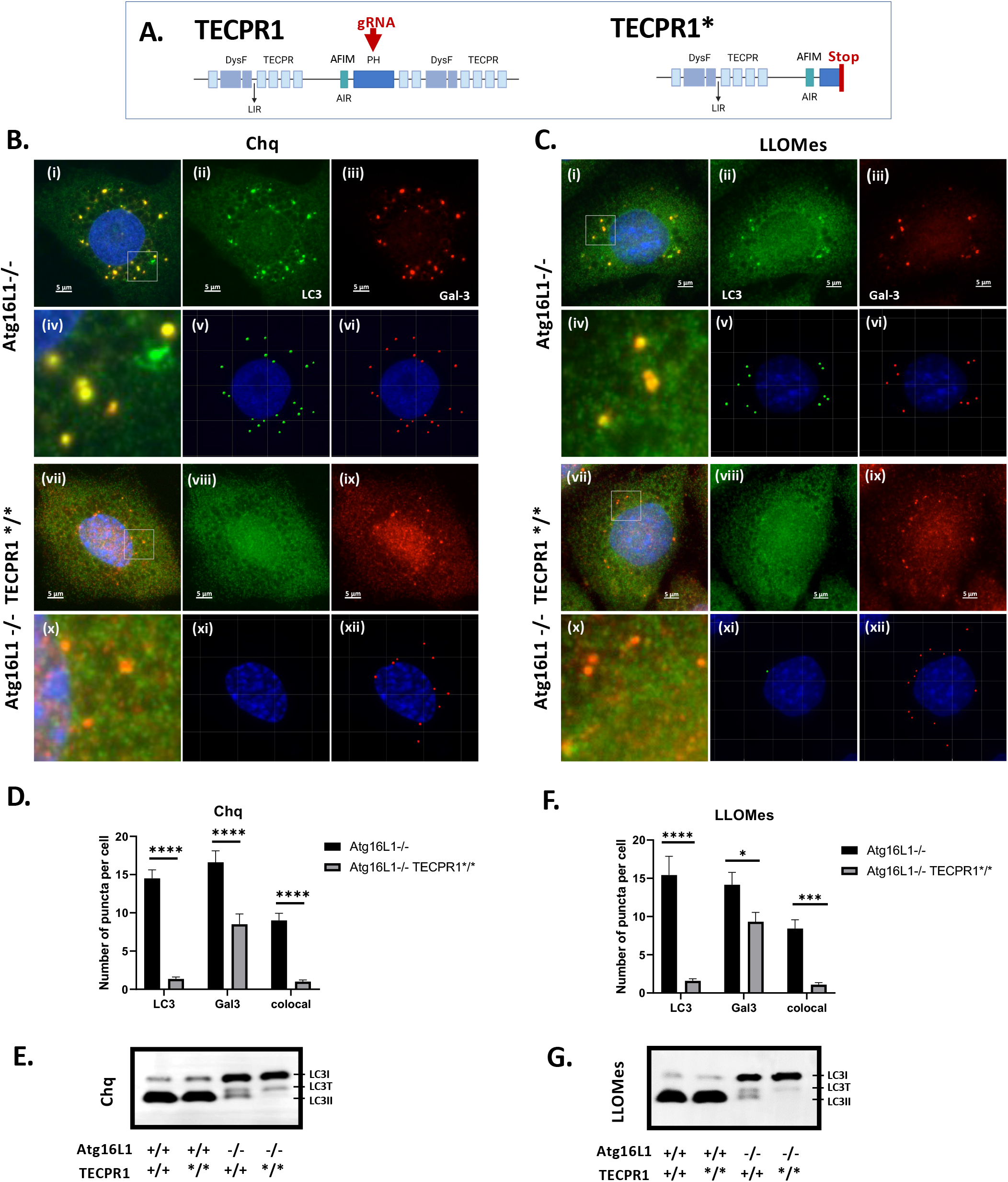
ATG16L1-independent conjugation of LC3 to damaged lysosomes requires TECPR1. **Panel A.** Domain map of TECPR-1 showing site of insertion of stop codon by custom CRISPR gRNA. Atg16L1-/-MEFs and gene edited Atg16L1-/-MEFS expressing truncated TECPR1* were incubated in nutrient media with chloroquine (100mM, **panel B**) or LLOMes (1mM **panel C**). Cells were fixed and immunostained for LC3(green) and galectin-3 (red). Rendered images (v and vi, xi and xii) were used to count puncta. Higher magnification images of boxed ROI are shown in iv and x. The graphs compare numbers of LC3 puncta co-localised with galectin-3 counted from rendered images of 20 cells incubated with chloroquine, P-values were calculated using multiple t-test (***P < 0.001). (**panel D**) or LLOMes (**panel F**). Generation of LC3II was assessed by western blot for LC3 (chloroquine **panel E**), (LLOMes **panel G**).

### TECPR-1 recruits LC3 and ATG5 to electron-dense membranes lacking the V-ATPase

TECRP-1 has an ATG5 interacting motif (AFIM) that can bind to the ATG5-ATG12 complex. The ability of TECPR-1 to recruit ATG5-ATG12 onto damaged lysosomes was tested by immunostaining for ATG5. Figure 5A shows that control MEFS generate puncta positive for galectin-3 and ATG5 following incubation with chloroquine (i-ii). Puncta labelling for galectin-3 and ATG5 were also produced in MEFS lacking ATG16L1 (iii-iv), but these were generated rarely in Atg16L1-/-MEFS expressing the truncated TECPR1 (v-vi). Similar results were obtained for co-staining for ATG5 and LC3 (supplemental data 2). Small bright puncta greater than 0.8mm diameter were counted from 10 cells selected at random. Quantification (supplemental data 2) confirms that recruitment of ATG5 to puncta positive for galectin-3 or LC3 in ATG16L1-/-MEFs was greatly reduced following truncation of TECPR1. The results provided further evidence that TECPR-1 recruits ATG5-ATG12 to conjugate LC3 to damaged lysosomes. CASM/LAP pathways induced by raised pH result in translocation of the V1 subunits of the V-ATPase to the cytoplasmic face of the vacuole where they recruit the ATG16L1:ATG5-ATG12 complex required for conjugation of LC3 to PE or PS [16, 30]. The distribution of the V-ATPase in cells incubated with chloroquine (Figure 5B) was used as a further means of distinguishing between swollen vacuoles and damaged lysosomes. As reported above, in control cells (Figure 5B i-iv). LC3 was recruited to the entire surface of the vacuole and to dense puncta close to the vacuoles in control cells. The V-ATPase was present on the limiting membrane of swollen vacuoles (large arrows) consistent with recruitment of LC3 via CASM/LAP, but the V-ATPase was absent from the bright LC3 puncta (small arrows). Similarly, the V-ATPase was recruited to the swollen vacuoles in ATG16L1-/-MEFs but not to the small LC3 puncta (Figure 5C i-iv). The results again suggested that recruitment of LC3 in the absence of ATG16L1 uses a pathway separate from pH dependent activation of the V-ATPase axis during LAP and CASM.

**Figure 5.**
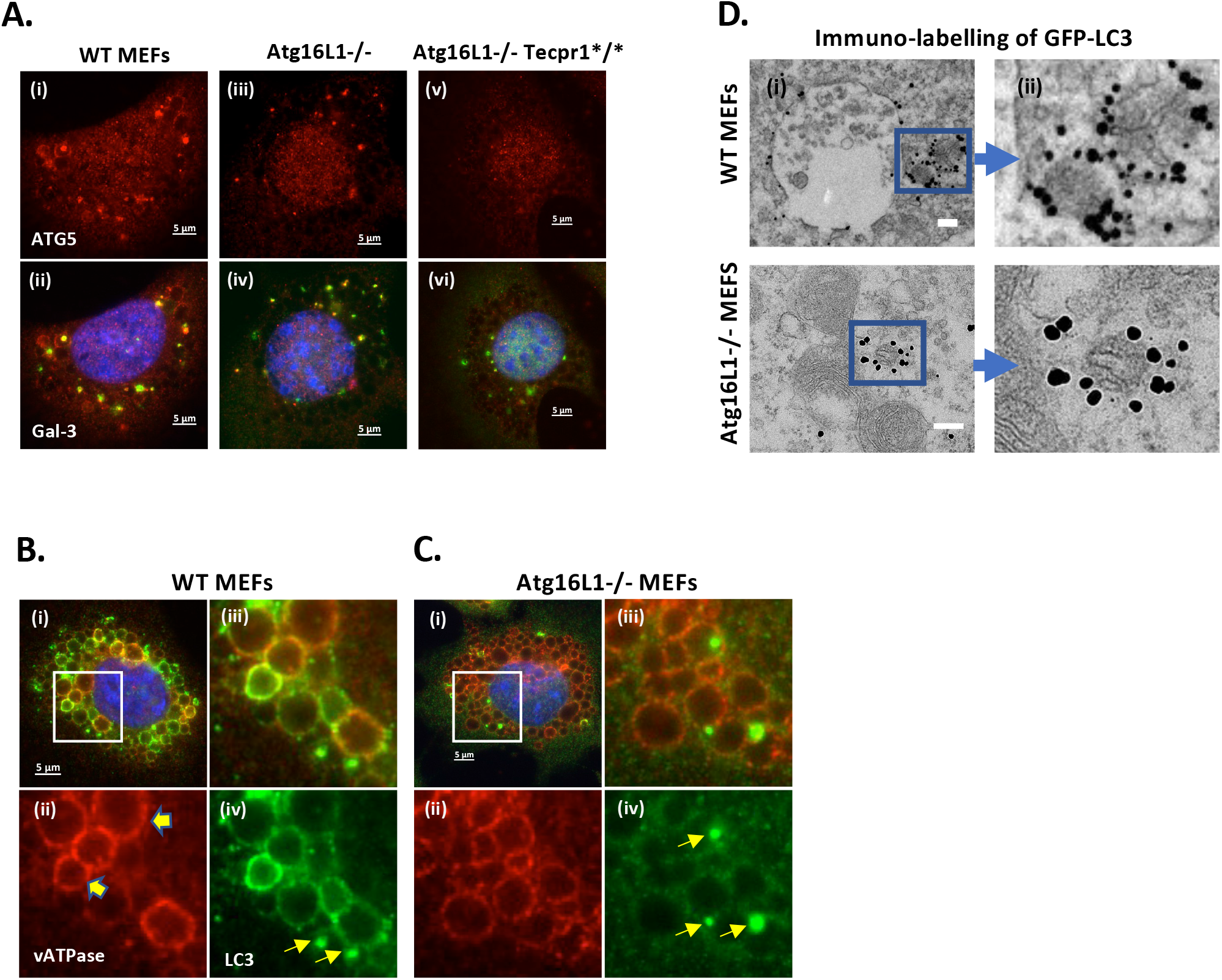
TECPR-1 recruits LC3 to electron dense membraneous material lacking the V-ATPase. Control (**panel A**) and Atg16L1-/-MEFs (**panel B**) were incubated for 2 hours in nutrient media containing chloroquine (100mM). Cells were fixed and immunostained for LC3 (green) and the V-ATPase (red). Regions of interest are shown at higher magnification (iii-iv). Large arrows indicate co-localised LC3 and V-ATPase on swollen vacuoles. Small arrows indicate small LC3 puncta that are negative for the V-ATPase. Panel C. Control (i-v) and Atg16L1-/-MEFs (vi-x) were transduced by adenovirus expressing GFP-LC3. Cells were incubated with chloroquine (100mM) for two hours and processed for immunogold electron microscopy. LC3 was visualised using antibody against GFP coupled to gold beads (1.4 nm), scale bar = 200nm.

The ultrastructure of the small bright LC3 puncta was analysed in MEFs transduced using an adenovirus expressing GFP-LC3. The location of LC3 puncta following incubation with chloroquine was determined by immunogold labelling for GFP-LC3 and searching for strong immunogold labelling close to swollen vacuoles (Figure 5D). In control cells the GFP-LC3 was largely localised to the limiting membrane of large (>1 µm diameter) swollen vacuoles (Fig 5D i) and to smaller electron dense (100-500nm) structures containing membranous material close by (Figure 5D ii). In ATG16L1-/-MEFs the strong immunogold labelling was again associated with small electron dense structures containing packed membranes (Figure 5D iii-iv). Recent studies [38] show that lysosome damage in response to LLOMes results in release of lysosome membrane proteins such as LAMP2 into the cytosol where they form small vesicles/complexes (also called lysosomal membrane complexes) that become a substrate for autophagy. The small diameter of 50-100nm of these structures and their content of LAMP2 resembles the electron dense structures reported here. This suggests that lysosomal membrane complexes released from damaged lysosomes may be the target for TECPR1 mediated conjugation of LC3.

### Role played by the LIR motif, PH domain and membrane association of TECPR1 during conjugation of LC3 to damaged lysosomes

TECPR1 tethers lysosomes to autophagosomes using an N-terminal tectonin repeat domain (TECRP) and a central PH domain to bind lysosomes while an LIR within the N-terminal TECPR domain binds LC3 on autophagosomes [35]. The role played by the LIR in recruiting LC3 to damaged lysosomes was tested by reconstituting ATG16L1-/-cells expressing the inactive truncated TECPR1* with full length TECPR1-RFP, or a mutant protein lacking residues W175A/I178A in the LIR required for LC3 binding (Figure 6). Expression of full length TECPR-1 reconstituted formation of LC3 puncta in response to chloroquine and the puncta were positive for TECPR-1 (Figure 6A i-iii) and galectin 3 (Figure 6B i-iii). This confirmed that TECPR-1 could locate to damaged lysosomes and recruit LC3 independently of ATG16L1. Conjugation of LC3 to lipids was confirmed by the appearance of LC3II on western blot (Figure 6C&D). Expression of the TECPR1-DLIR (W175A/I178A) mutant failed to restore LC3 puncta (Figure 6A iv-vi) and TECPR1-DLIR was absent from damaged lysosomes positive for galectin-3 (Figure 6B iv-vi) and LC3II was absent from western blots (Fig 6C&D). Interestingly, the TECPR1-DLIR mutant was not recruited to galectin-3 puncta (Figure 6B iv-vi). Given that the W175A/I178A residues lie within the TR1 domain required for lysosome binding it is possible that the W175A/I178A mutation inactivates lysosome binding properties of the TR1 domain and recruitment of TECPR-1 to damaged lysosomes and conjugation of LC3 are no longer possible. Figure 3 has shown that damaged lysosomes generated by chloroquine or LLOMes are enriched for PI4P. The role played by the PI4P-specific PH domain of TECPR1 in recruiting LC3 to damaged lysosomes was therefore tested by repeating the experiment using TECPR1 lacking the PH domain (TECPR1-DPH). LC3 puncta were restored following expression of TECPR1-DPH RFP (Figure 6A vi-ix) and these co-located with TECPR1-DPH RFP, but numbers of puncta were lower than seen for full length TECPR-1. The TECPR1-DPH RFP mutant located to puncta, and some co-stained for galectin 3 (Figure 6B vi-ix) while others were negative for galectin or LC3. This was also evident in western blots showing low levels of LC3II generated in response to chloroquine. This suggested that recruitment of LC3 to damaged lysosomes was reduced following loss of the PH domain, but not lost completely. It is possible that binding to lysosomes is maintained by the TRI domain that remains intact in the TECPR1-DPH mutant.

**Figure 6.**
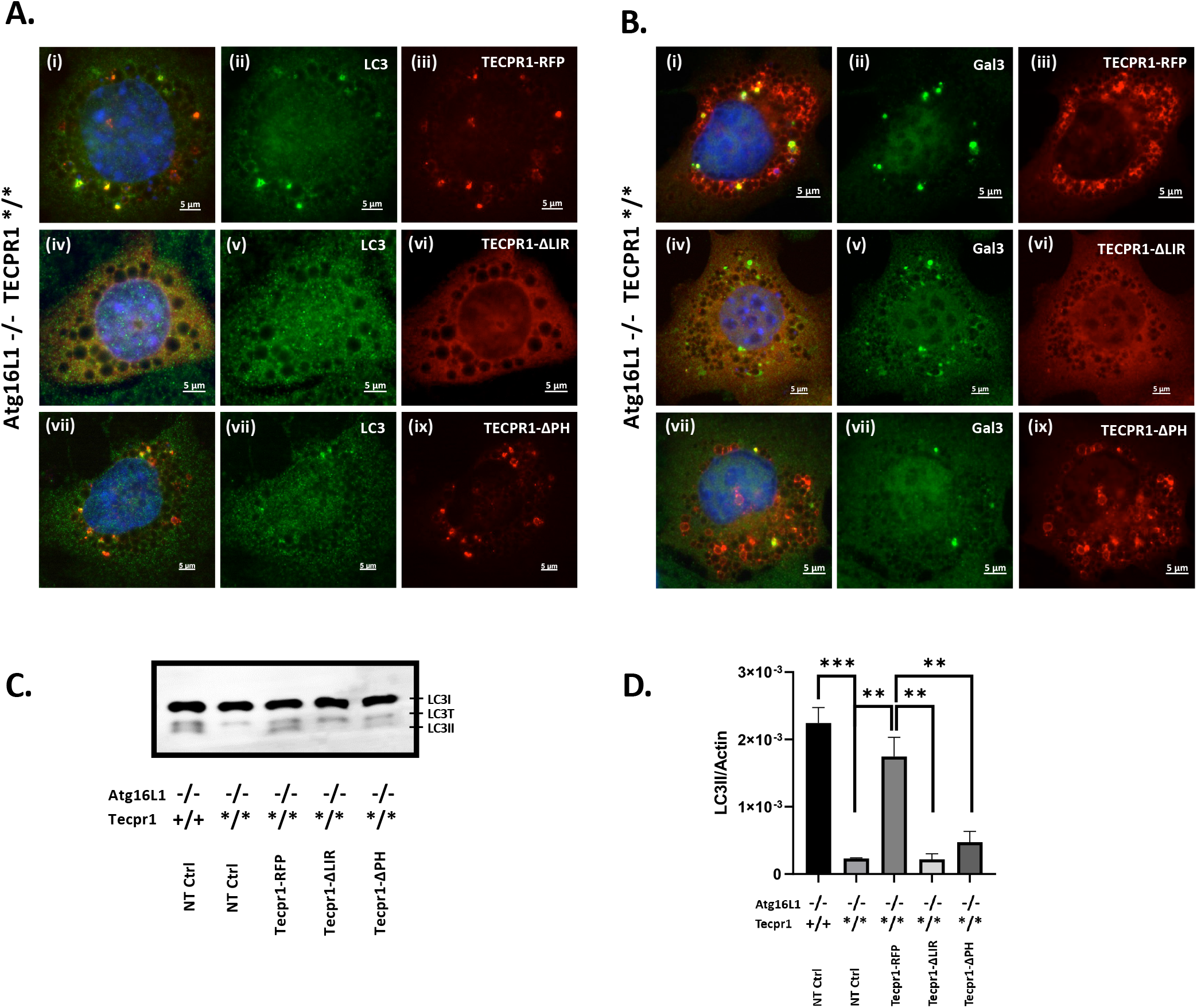
Role played by the LIR motif, PH domain and membrane association of TECPR-1during conjugation of LC3 to damaged lysosomes. Gene edited Atg16L1-/-MEFS expressing truncated TECPR1* were transfected with plasmids expressing full length TECPR-1 RFP (i-iii), TECPR-1 lacking the LIR within the N-terminal TECPR domain (Tecpr1-W175A/I178A-GFP pseudo red, iv-vi) or TECPR-1 lacking the central PH domain (Tecpr1-dPH-GFP pseudo red, vii-ix). Cells were incubated in nutrient media containing chloroquine (100µM) for 2 hours. **Panel A** shows counter staining for LC3 (green) and **panel B** counter staining for galectin 3 (green). **Panel C and D**. Western blot analysis and relative quantification of LC3II. Atg16L1-/-TECPR*/*MEFS transfected with indicated plasmids were incubated in nutrient media containing chloroquine (100µM) for 2 hours. Data represent mean ± S.E. of three independent experiments (***P < 0.001).

## Discussion

The possibility that proteins other than ATG16L1 could provide E3 ligase activity to the ATG5-ATG12 complex was tested by incubating cells lacking ATG16L1 under conditions known to activate autophagy. Activation of canonical autophagy by starvation did not lead to the formation of LC3 puncta or LC3II. In contrast bright LC3 puncta and LC3II were produced when ATG16L1-/-cells were incubated with chloroquine to raise vacuolar pH and activate the CASM/LAP/V-ATPase pathway. In control cells, LC3 was recruited to the entire limiting membrane of swollen vacuoles, indicating recruitment via CASM/LAP, and to smaller bright puncta close to the swollen vacuoles. In contrast, when ATG16L1 was absent, LC3 was only recruited to the small bright puncta, even though the swollen vacuoles were positive for the V-ATPase. This localised accumulation of LC3 in bright puncta rather than the vacuole membrane, and the lack of requirement for the WD domain of ATG16L1, suggested that recruitment of LC3 to the small bright puncta was separate from the CASM/LAP/V-ATPase pathway. Importantly, conjugation of LC3 in the absence of ATG16L1 still required proteins upstream of ATG16L1 such as ATG3, ATG5 and ATG7, and ATG5 was recruited to sites of LC3 conjugation. The ATG16L1-independent conjugation of LC3 to the small bright puncta in response to chloroquine therefore required the E1 and E2 -ubiquitin-like activities of ATG7 and ATG3 that activate LC3 prior to processing by the E3 ligase-like activity of the ATG5-ATG12 conjugate. Further analysis showed that the bright LC3 puncta shared many characteristics with damaged lysosomes. The puncta were positive for LAMP and exposed luminal galactoside sugars to the cytosol where they were recognised by galectin-3. Identical puncta were formed by the lysosome damaging compound LLOMes which also increased levels of PI4P in the LC3 positive puncta suggesting activation of lysosome repair pathways and recruitment of phosphatidylinositol 4-kinase[37]. Ultrastructural analysis revealed that the bright LC3 puncta were electron dense structures packed with internal membranes. These were similar to the lysosomal membrane complexes released into the cytosol following permeabilization of lysosomes by LLOMes [38].

The observations that TECPR1 binds ATG5 [31, 32] and that cells generate separate ATG16L1 or TECPR1 complexes with ATG5-ATG12 [34] suggested that TECPR1 could substitute for ATG16L1 during conjugation of LC3 to membranes. This was supported by the observation that LC3 conjugation in response to chloroquine or LLOMes was prevented by insertion of a stop codon into exon 12 of the central PH domain of TECPR1, even though galectin-3 staining showed that lysosomes were damaged. Subsequent reconstitution experiments showed that expression of full length TECPR1 could restore formation of LC3 puncta and LC3II in ATG16L1-/-MEFs expressing the truncated inactive TECPR1*. This provided strong evidence that TECPR1:ATG5-ATG12 complexes conjugate LC3 to damaged lysosomes. By analogy with ATG16L1, it is possible that ATG5 in the TECRP1:ATG5-ATG12 complex induces a conformational change in ATG3 to transfer the C-terminal carboxyl group of LC3 from a thioester bond to amino groups exposed by PE. The site of LC3 lipidation during canonical autophagy is determined by the location of ATG16L1 [26, 28]. The coiled coil domain of ATG16L1 binds WIPI which in turn binds PIP3 to direct LC3 conjugation to sites of autophagosome formation. During LAP/CASM the WD domain of ATG16L1 binds the V-ATPase to direct LC3 lipidation to swollen vacuoles in response to raised pH [16]. The ability of TECPR1 to bind PI4P may play a role by directing TECPR1 to sites of lysosome damage and repair but loss of the PH domain only had a partial effect suggesting further signals are required to activate LC3 conjugation. Conjugation of LC3 to damaged lysosomes was dependent on the N-terminal LIR suggesting a role for LC3 binding, however the modified W175A/I178A residues of the LIR lie within the TECPR domain required for lysosome binding. It is not possible therefore to exclude the possibility that the W175A/I178A mutation inactivates lysosome binding properties of the N-terminal TECPR domain making recruitment of TECPR1 to damaged lysosomes and conjugation of LC3 no longer possible.

The ability of TECPR1 to conjugate LC3 directly to lysosomes may contribute to the tethering functions of TECPR1 reported to promote autophagosome lysosome fusion [33, 35] during the removal of protein aggregates. Tethering is achieved through the N-terminal TECPR domain that binds lysosomes, and an LIR able to bind LC3 on autophagosomes, and this may be enhanced by the central PH domain in response to the production of PI4P during lysosome damage. An accumulation of protein aggregates in autophagosomes and/or lysosomes may damage vacuoles exposing signals that recruit TECPR1 to tag the vacuoles with LC3 for removal or repair. TECPR1 is recruited to bacterial targets for autophagy such as *ΔicsB Shigella, S. typhimurium*, Group A Streptococcus and to depolarised mitochondria [32]. Knock down of TECPR1 reduces recruitment of LC3 to bacteria and increases bacterial replication in cells suggesting that TECPR1 promotes degradation of bacteria by autophagy. Similarly, MEFs from TECPR1-/-mice show defects in removal of protein aggregates and disposal of depolarised mitochondria implicating roles for TECPR1 in aggregate clearance and mitophagy [32]. It will be interesting to determine if TECPR1 can conjugate LC3 to bacteria during xenophagy independently of ATG16L1 as a defence against infection. This is possible because in common with damaged lysosomes, vacuoles containing bacteria expose luminal sugars to the cytosol and recruit galectin-3 [39].

The observation that TECPR1 binds WIPI-2 [32] provides further similarity with ATG16L1, however unlike ATG16L1, TECPR1 does not contribute to early stages in autophagosome formation but is required later during autophagosome maturation [33]. In common with WIPI-2, TECPR1 has a PH domain able to bind PI3P but the interaction of TECPR1 with PI3P may be subject to autoinhibition. Studies ‘in vitro’ show that the PI3P binding is inhibited by the neighbouring AIM domain and that this autoinhibition is relieved when TECPR1 binds ATG5-ATG12 [33]. It will be interesting to see if AIM-dependent autoinhibition also regulates binding of TECPR1 to damaged lysosomes.

The repair and/or removal of damaged lysosomes promotes cell survival by preventing leakage of H^+^ and hydrolytic enzymes into the cytosol. Soon after initiation of damage ESCRIT-III proteins accumulate on damaged lysosome where they locate to subdomains of the damaged vacuole [40, 41] and may package damaged regions for exocytosis, or make repairs in situ, as they do at the plasma membrane and nuclear envelope. ESCRIT recruitment occurs before LC3 recruitment and repair prevents lysophagy. During lysophagy, recruitment of galectin-8 to damaged lysosomes inhibits the Ragulator-Rag amino acid sensor in the lysosome membrane leading to inhibition of mTOR and activation of autophagy [42]. At the same time ubiquitination of lysosome content and exposure of gaectin-3 leads to recruitment of cargo receptors (p62, OPTN, TAX1BP1) that bind to LC3 in the autophagosome membrane resulting in engulfment and fusion with undamaged lysosomes. It is possible that TECPR1-dependent conjugation of LC3 to damaged lysosomes increases lysophagy by increasing the availability of LC3 for recruitment of cargo receptors, tethering proteins (HOPS, PLEKHM1) and SNARE proteins (STX17, SNAP29, VAMP8) required for fusion with lysosomes.

Removal by lysophagy is counter balanced by autophagic lysosomal reformation (ALR) pathways activated by lysosomal damage. During ALR tubular structures extend from the lysosome and release small vesicles called proto-lysosomes containing lysosomal membrane proteins but devoid of luminal proteases [43]. Tubulation is driven by changes in phospholipid (PI94,5)P_2_, PI4P, PI3P) composition in microdomains of the lysosome. These domains recruit the kinesin microtubule motor protein KIF5B which drives tubulation of the lysosome and scission of the tubules by dynamin-2 allowing the released proto-lysosomes to recycle lysosomal membrane proteins to repair and regenerate lysosomes. Recent LC3 interactome analysis shows that lysosome damage induces binding of ATG8/LC3 to lysosome membrane protein LIMP2 [44]. LC3 also recruits TBC1D15, a Rab7 GTPase activating protein that activates Rab7 to promote dynamim-2 dependent release of proto-lysosomes. It is possible that conjugation of LC3 to the surface of the damaged lysosome by TECPR1 can facilitate recruitment of TBC1D15 to increase release of proto-lysosomes dynamin-2 making them available for lysosome repair.

The lysosome damage response also activates TFEB to promote transcription of genes important during autophagy and lysosome biogenesis. During this process leakage of Ca^2+^ from lysosomes activates autophagy leading to conjugation of LC3 to the lysosome membrane where it binds the TRPML1 calcium channel to activate TFEB [45]. It will be interesting to determine if TECPR1-mediated conjugation of LC3 to damaged lysosomes contributes to TFEB activation in response to damage. The galectin-3/LC3 positive puncta generated by TECPR1 in ATG16L1-/-cells may represent intermediates in the lysosome repair/removal pathways because the cells are unable to activate downstream pathways requiring ATG16L1. One example involves the binding of TRIM16 to galectin-3 in response to lysosome damage [46]. This allows TRIM16 to bind ATG16L1 and influence TFEB and mTOR during lysophagy and phagosome repair. These arms of the galectin-TRIM16 axis would be lost in cells lacking ATG16L1.

## Materials and Methods

### Antibodies and reagents

rabbit-LC3A/B (CST #4108), mouse-LC3B (CST #83506), rat-Lamp1 (abcam [1D4B] ab25245), rat-galectin3 (Santa Cruz M3/38 sc-23938), rabbit-Atg5 (GTX113309), GP-p62 (Progen GP62-C), mouse-EEA1 (MBL M176-3), mouse-Actin (sigma A2228). Chloroquine (Chq) (Sigma, C-6628, 100µM), monensin (Mon) (Sigma, M-5273, 100µM), Leu-Leu methyl ester hydrobromide (LLOMes) (Sigma, L-7393, 1mM), bafilomycin A (Baf) (Sigma, B-1793, 100nM), diphenyleneiodonium chloride (DPI) (Sigma, D-2926, 10µM), phloretin (phl) (Sigma, P-7912, 180µM), wortmannin (Wtm) (Sigma, W-1628, 100nM)

### Plasmid and transfection

Cells were transfected with FuGENE® HD Transfection kit according to the manufacturer’s instructions, harvested or analysed 24–48 hours post transfection. Tecpr1-RFP, Tecpr1-W175A/I178A-GFP, Tecpr1-dPH-GFP were generous gifts from Stephane Blanchard and Thomas Wollert and described in Wetzel et al 2020.

### Cells and cell culture

Mouse embryonic fibroblasts (MEFs) were cultured in DMEM (ThermoFisher scientific, 11,570,586) with 10% fetal bovine serum and 2mM L-glutamine and 100U/ml penicillin/streptomycin and incubated in 5% CO2 at 37°C. Cells were starved by incubating in Hank’s balanced salt solution (HBSS; Invitrogen, 14025-076) which lacks amino acids for the specified length of time to stimulate autophagy.

### Truncation of Tecpr1 in MEFS

Gene editing of Tecpr1 in MEFs was achieved using custom CRISPR gRNA lentivirus transduction particles (Mission, Sigma Aldrich). MEFs were infected with the CRISPR gRNA lentiviruses in Opti-Minimal Essential Medium (Opti-MEM) at MOI of 1, supplemented by 16 μg/mL hexadimethrine bromide. Transduced cells were selected by 10 μg/mL puromycin and single cell colonies were isolated by dilution. Genomic regions containing the editing site were amplified by PCR and screened for frameshift mutations by sequencing. (gRNAs 5’-AGCAGTTCACAGGGCACGA and gRNA 5’-TGTACACGGGCGGCTATGG)

### Western blotting

Protein was extracted using M-PER (ThermoFisher Scientific, 78,501) with complete protease inhibitor cocktail (Sigma, 04693159001) for 30 min on ice. Samples were clarified by centrifugation (13,200g, 7 min). From the supernatants, protein concentrations were determined using the BCA protein assay system (ThermoFisher Scientific, 23,225). Protein (30 µg) was separated on a precast 4–12% gradient or 12% np gradient SDS-PAGE gel (Expedeon, NBT41212) and transferred to immobilon PVDF (Millipore, IPFL00010) for blotting. Membranes were probed using antibodies for ATG16L1 (MBL M150-3) and ACTB/actin (Sigma, A5441). Primary antibodies were detected using IRDye labelled secondary antibodies (LI-COR biosciences, 926–32,211, 926–68,020) at 1:10,000 dilution. Proteins were visualized using the Odyssey infrared system (LI-COR).

### Fluorescence microscopy

Cells grown on glass coverslips in 24-well plates were fixed at −20°C in ice cold methanol for 7 min, then blocked in 5% goat serum, 0.3% Triton X-100 in PBS (Sigma, G9023; X100) for 30 mins. Cells were incubated with primary antibody overnight. Washed cells were incubated with secondary antibody anti-Rabbit IgG Alexa Fluor® 488 (Life Technologies, A11008), anti-Rat IgG Alexa Fluor®594, anti-Rat IgG Alexa Fluor®488 and anti-mouse IgG Alexa Fluor® 594, 2hrs at room temperature, followed by counterstain with 4′, 6 diamidino-2-phenylindole (DAPI; ThermoFisher Scientific, 10,116,287) and mounted on slides with Fluoromount-G from Southern biotech (ThermoFisher Scientific, 15,586,276). Cells were imaged on a Zeiss Imager M2 Apotome microscope with a 63X, 1.4 NA oil-immersion objective using 365 ± 40 nm excitation and 445 ± 25 nm emission for DAPI, 470 ± 20 nm excitation and 525 ± 25 nm emission for LC3.

## Acknowledgements

We are very grateful to Thomas Wollert and Stephane Blanchard for providing plasmids expressing TECRP1-RFP, Tecpr1-W175A/I178A-GFP and Tecpr1-DPH-GFP [35] and to Norwich Medical School, University of East Anglia for support of PhD studies by KK.

## Figure legends

**Supplemental figure 1. Crispr/Cas9 gene editing of Tecpr1 using custom CRISPR gRNA lentivirus transduction particles**. A sequence schematic highlighting the sequence of the gRNA and location of the targeted site in the Tecpr1 (**panel A**). Sequence alignments of the targeted region in selected mutant (**panel B**). An insertion has been detected in the gRNA sequence, which induces a frameshift and a premature stop codon within exon 12 of the PH domain of Tecpr1, resulting in a truncated Tecpr1. **(panel C)**. Sanger sequencing results **(panel D)**.

**Supplemental figure 2. TECPR-1 recruits LC3 and ATG5 to damaged lysosomes**. Control, Atg16L1-/- and ATG16L1-/-MEFS expressing truncated TECPR1* were incubated for 2 hours in nutrient media containing chloroquine (100μM). **Panel A**. Cells were fixed and immunostained for ATG5 (red) and galectin-3 (green). **Panel B**. Cells were fixed and immunostained for ATG5 (red) and LC3 (green). **Panel C**. Quantification and composition of puncta positive for ATG5 and galectin 3. **Panel D**. Quantification and composition of puncta positive for ATG5 and LC3. n ≥ 10 cells were quantified by imaris, P-values were calculated using multiple t-test (***P < 0.001).

