## Supplemental figures for "TECPR1 provides E3-ligase like activity to the ATG5-ATG12 complex to conjugate LC3/ATG8 to damaged lysosomes"

### Supplemental figure 1:

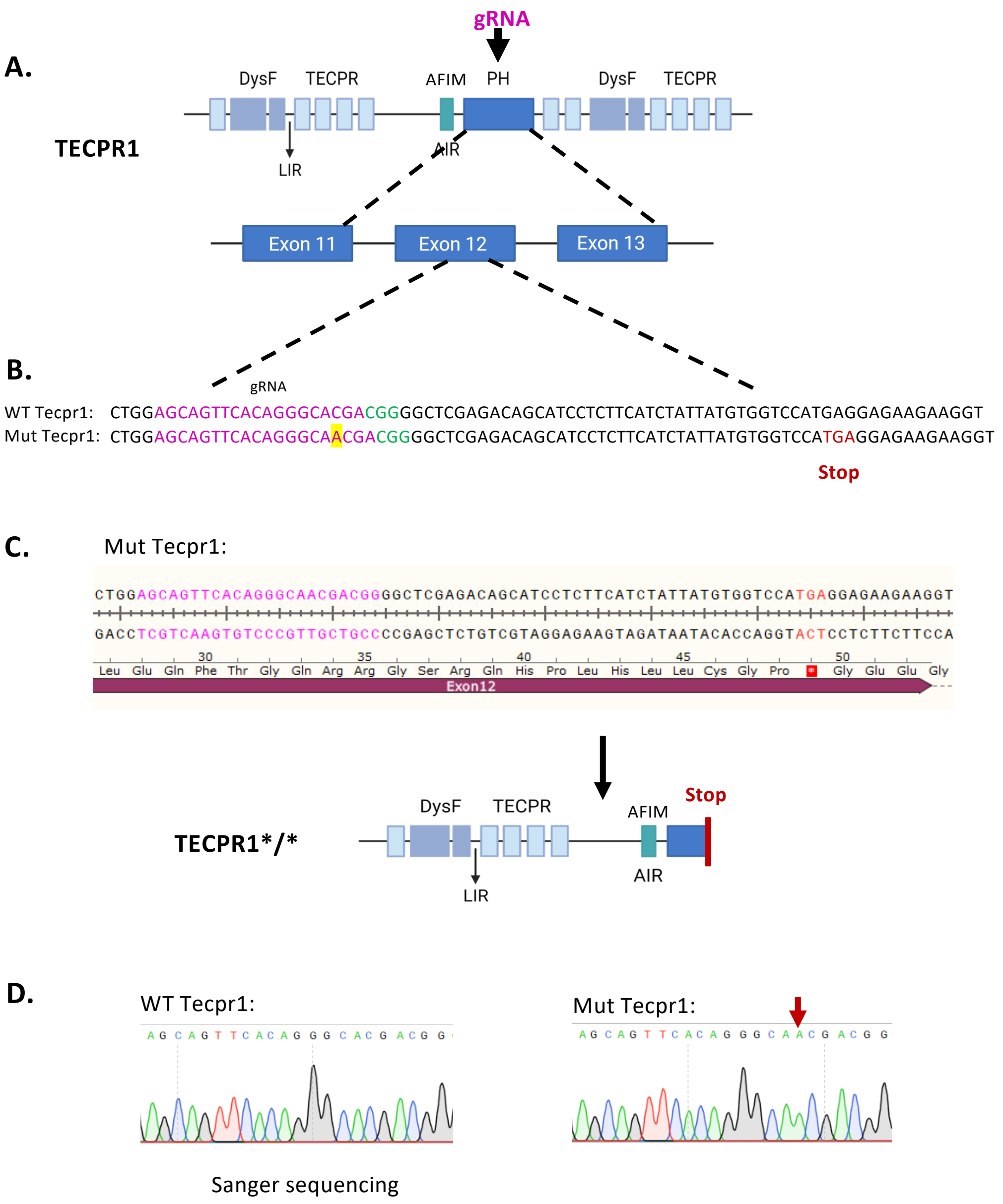


**Supplemental figure 1. Crispr/Cas9 gene editing of Tecpr1 using custom CRISPR gRNA lentivirus transduction particles**. A sequence schematic highlighting the sequence of the gRNA and location of the targeted site in the Tecpr1 (**panel A**). Sequence alignments of the targeted region in selected mutant (**panel B**). An insertion has been detected in the gRNA sequence, which induces a frameshift and a premature stop codon within exon 12 of the PH domain of Tecpr1, resulting in a truncated Tecpr1. **(panel C).** Sanger sequencing results **(panel D)**.

**Supplemental figure 2:**

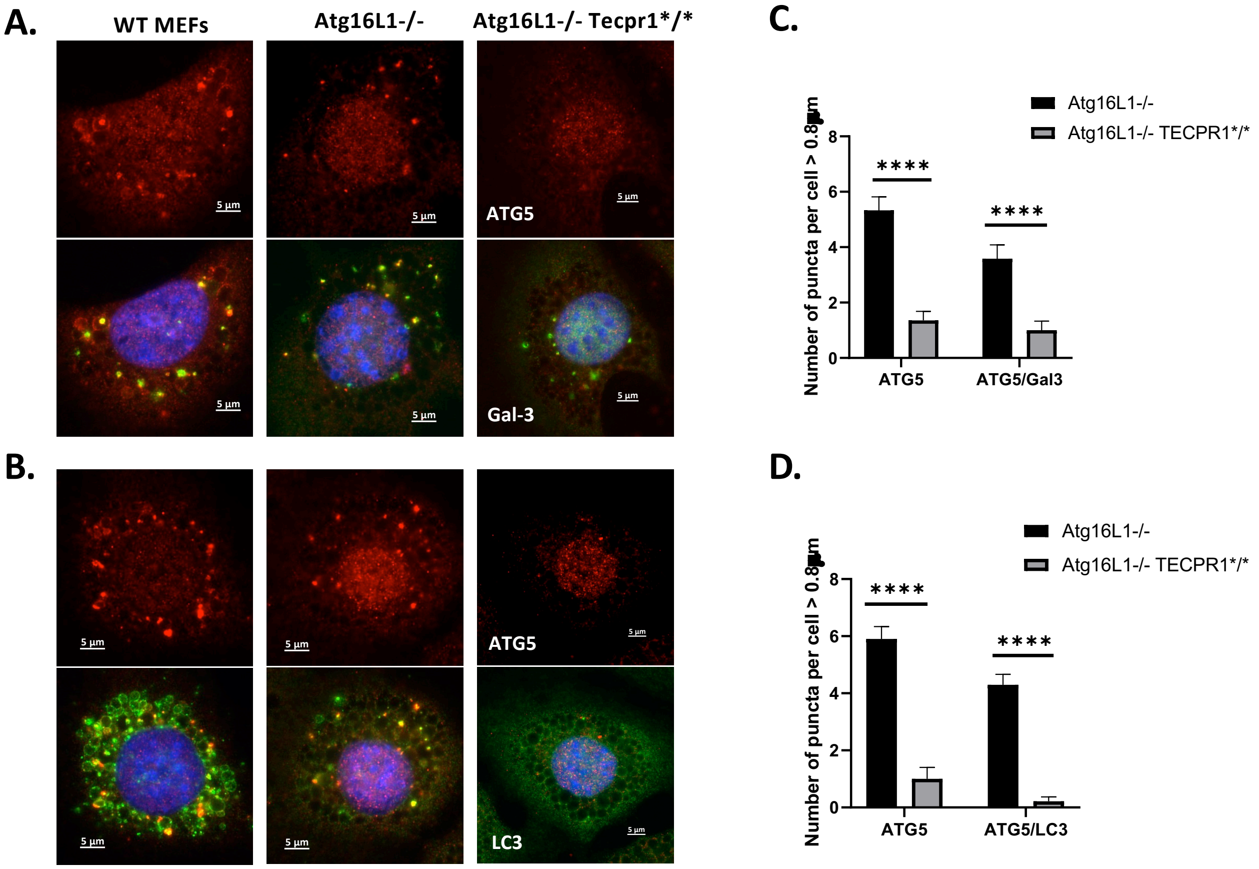


**Supplemental figure 2. TECPR-1 recruits LC3 and ATG5 to damaged lysosomes.** Control, Atg16L1-/- and ATG16L1-/- MEFS expressing truncated TECPR1* were incubated for 2 hours in nutrient media containing chloroquine (100μM). **Panel A.** Cells were fixed and immunostained for ATG5 (red) and galectin-3 (green). **Panel B.** Cells were fixed and immunostained for ATG5 (red) and LC3 (green). **Panel C.** Quantification and composition of puncta positive for ATG5 and galectin 3. **Panel D.** Quantification and composition of puncta positive for ATG5 and LC3. n ≥ 10 cells were quantified by imaris, P-values were calculated using multiple t-test (***P < 0.001).
